# Pretrained language models and weight redistribution achieve precise *k*_cat_ prediction

**DOI:** 10.1101/2022.11.23.517595

**Authors:** Han Yu, Xiaozhou Luo

## Abstract

The enzyme turnover number (*k*_cat_) is a meaningful and valuable kinetic parameter, reflecting the catalytic efficiency of an enzyme to a specific substrate, which determines the global proteome allocation, metabolic fluxes and cell growth. Here, we present a precise *k*_cat_ prediction model (PreKcat) leveraging pretrained language models and a weight redistribution strategy. PreKcat significantly outperforms the previous *k*_cat_ prediction method in terms of various evaluation metrics. We also confirmed the ability of PreKcat to discriminate enzymes of different metabolic contexts and different types. Additionally, the proposed weight redistribution strategies effectively reduce the prediction error of high *k*_cat_ values and capture minor effects of amino acid substitutions on two crucial enzymes of the naringenin synthetic pathway, leading to obvious distinctions. Overall, the presented *k*_cat_ prediction model provides a valuable tool for deciphering the mechanisms of enzyme kinetics and enables novel insights into enzymology and biomedical applications.

## Introduction

Understanding enzyme catalysis efficiency to a specific substrate is an essential and fundamental biological problem that affects metabolism, metabolite concentrations and fluxes, and resource allocation in enzymes [1–4]. The value of *k*_cat_, the maximal turnover rate, reflects an enzyme’s catalytic activity to a large extent. Various databases, such as UniProt [5], Brenda [6], and SABIO-RK [7], have provided easy-to-access interfaces to acquire a large number of experimentally measured *k*_cat_ values. However, the scales of the *k*_cat_ values databases are far smaller than those of enzyme sequences. For instance, the number of enzyme sequences exceeds 230 million, while the number of experimentally measured *k*_cat_ values is just above 2000 in UniProt [5]. This large gap limits the development of downstream applications and needs to be overcome.

Currently, the measurement of the *k*_cat_ value mainly depends on experimental measurements, yet it is human-demanding, time-consuming, and expensive. Therefore, several studies have tried to utilize computational methods to accelerate the process of *k*_cat_ prediction. Simon verified that enzyme kinetic parameters could be predicted well based on statistical learning [8]. Heckmann demonstrated that machine learning could predict catalytic turnover numbers in Escherichia coli based on enzyme biochemistry, protein structure, and network context [9]. More representative, Feiran proposed deep learning-based *k*_cat_ prediction solely from substrate structures and protein sequences, which realized high-throughput prediction [10]. However, the model performance needs to be greatly improved. For instance, the coefficient of determination on the test dataset is only approximately 0.48 [10], indicating that there is still room for improvement. In addition, truly experimental cases rather than the datasets of the existing databases need to be tested for the validation of model robustness.

However, a novel data representation approach based on an unsupervised learning method originating from natural language processing has yielded great breakthroughs [11–12]. It has also been applied to biological problems [13–15]. For instance, protein representation learning based on a pretrained language model has been verified to be effective in various related tasks, such as protein secondary structure, protein subcellular localization, biochemical properties of amino acids and remote homology of proteins [14]. Similarly, substrate molecular representation utilizing a pretrained SMILES Transformer has confirmed that it facilitates molecular property prediction tasks [15]. The pretrained language models may facilitate *k*_cat_ prediction.

In addition, because the current model cannot tackle high value predictions well on an imbalanced dataset, which is usually given more attention, it is worth being optimized. To process imbalanced datasets, some studies have used various tricks to avoid them [16]. Recently, Yuzhe defined the concept of deep imbalanced regression and proposed label distribution smoothing and feature distribution smoothing to reduce the prediction error [17]. Another common method is to adjust the sample weight distribution, which has been verified to be very effective on related biological problems [18]. These methods could be applied to solve the high value prediction problem on an imbalanced dataset.

Here, we propose a pretrained language model-based Kcat prediction approach (PreKcat), which precisely predicts the *k*_cat_ value from the enzyme sequence and substrate structure. We demonstrated its remarkable superiority compared with the previous *k*_cat_ prediction method. Furthermore, we employed a weight redistribution strategy to improve high value prediction on an imbalanced dataset. The experimental results of two crucial enzymes on the naringenin synthetic pathway also proved the effectiveness of PreKcat on enzyme variant prediction.

## Results

### Overview of PreKcat

The framework of PreKcat is composed of a representation and a machine learning module, where the former is used to represent enzyme and substrate information based on pretrained language models, and the latter utilizes the representative machine learning method Extra Tree model to predict the corresponding *k*_cat_ value (Fig. 1). Specifically, each amino acid of the enzyme sequence was converted into a 1024-dimensional vector by the protT5-XL model, and we used mean pooling to benchmark the per-protein predictions, which has been verified to be most effective on per-protein tasks compared with other pooling methods [13]. The substrate structure is first converted into a simplified molecular-input line-entry system (SMILES) and fed into the pretrained SMILES transformer, which generates a 256-dimensional vector for each amino acid. The mean and max pools of the penultimate and last hidden layers then generate 256-dimensional vectors for each char of SMILES, respectively [15]. Then, these embedded vectors generate a per-molecular representation for a substrate. Furthermore, the concatenated representation vector of protein and substrate are imputed into the explainable extra tree model, which has been clarified to be useful on various protein or peptide function prediction tasks [19–20]. We also found that a solely concatenated representation vector cannot discriminate high or low *k*_cat_ values well by projection of t-distributed stochastic neighbour embedding (t-SNE) [21], further demonstrating the necessity of the machine learning model (Supplementary Fig. 1).

**Fig. 1.**
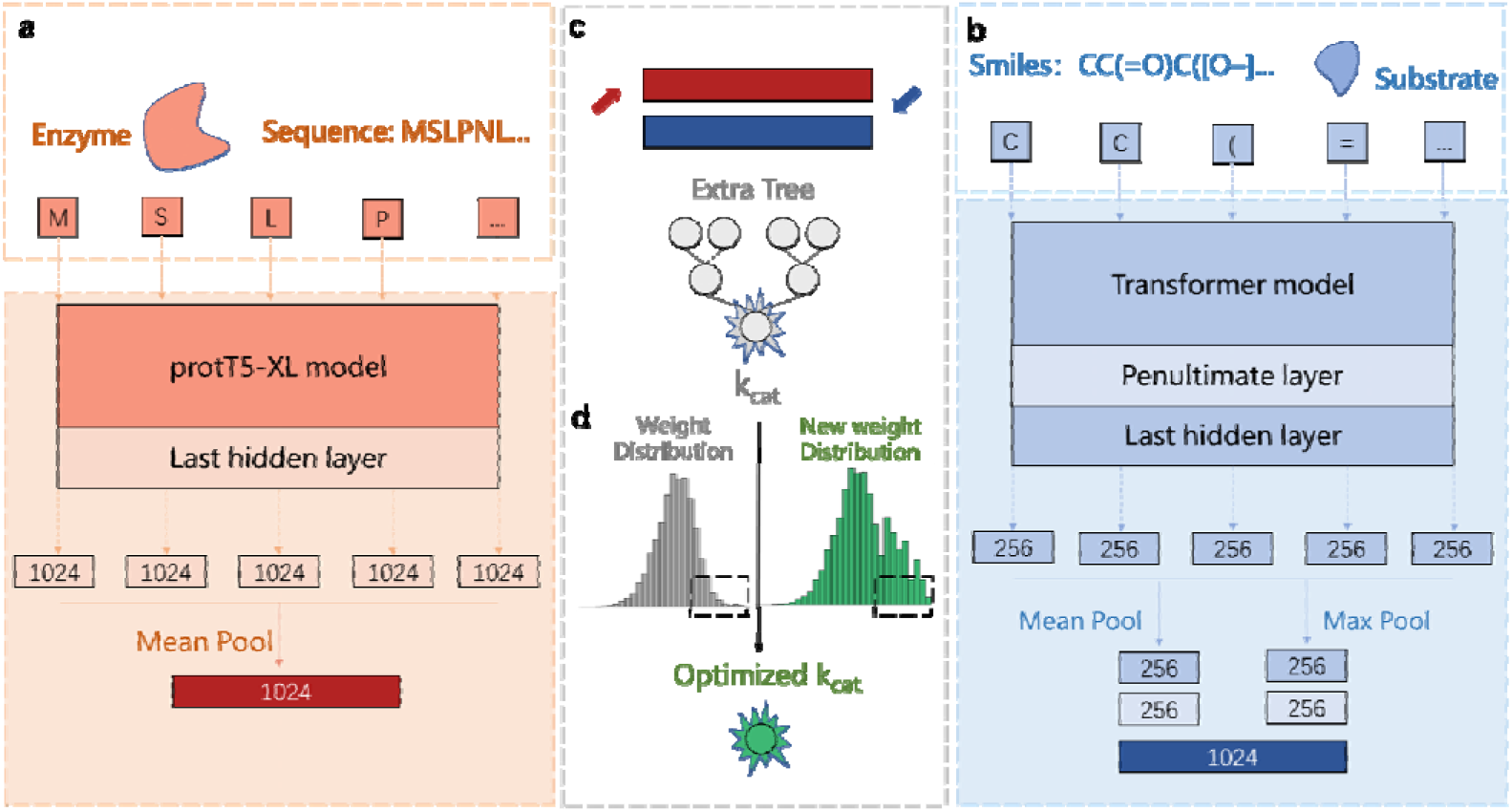
Overview of PreKcat. a, Enzyme sequence representation module. b, Substrate structure representation module. c, Machine learning module. d, Weight redistribution of the sample or feature.

### PreKcat significantly outperforms the previous method in *k*_cat_ prediction

To evaluate our proposed model, we selected the DLKcat dataset, which includes 16838 samples, and compared PreKcat with that dataset, referring to its evaluation metrics [10]. Without additional parameter optimization, the average value of the coefficient of determination on the test dataset of five rounds of random splitting was 0.66, which was 18% higher than that of DLKcat (Fig. 2a). This marked advantage demonstrated the superiority of PreKcat, and the smaller variance also verified its robustness. Meanwhile, the root mean square error (r.m.s.e.) values of the predicted *k*_cat_ values and experimentally measured *k*_cat_ values were also smaller than those of the previous model on the training and test sets (Fig. 2b). The mean absolute error (m.a.e.) was also relatively small (Supplementary Fig. 2). Moreover, we found that a high correlation between the two groups of *k*_cat_ values existed in the test set (Fig. 2c; Pearson correlation coefficient (p.c.c.) = 0.82) and the whole dataset (Supplementary Fig. 3; p.c.c. = 0.98) and were 11% and 10% higher than DLKcat, respectively. Additionally, PreKcat presented better prediction performance on the test set where at least either the enzyme or the substrate was not in the training set (Supplementary Fig. 4; p.c.c. = 0.80). The performance on various *k*_cat_ numerical intervals also demonstrated the superiority of PreKcat (Supplementary Fig. 5). Altogether, PreKcat significantly outperformed DLKcat in enzyme *k*_cat_ prediction in terms of various evaluation metrics.

**Fig. 2.**
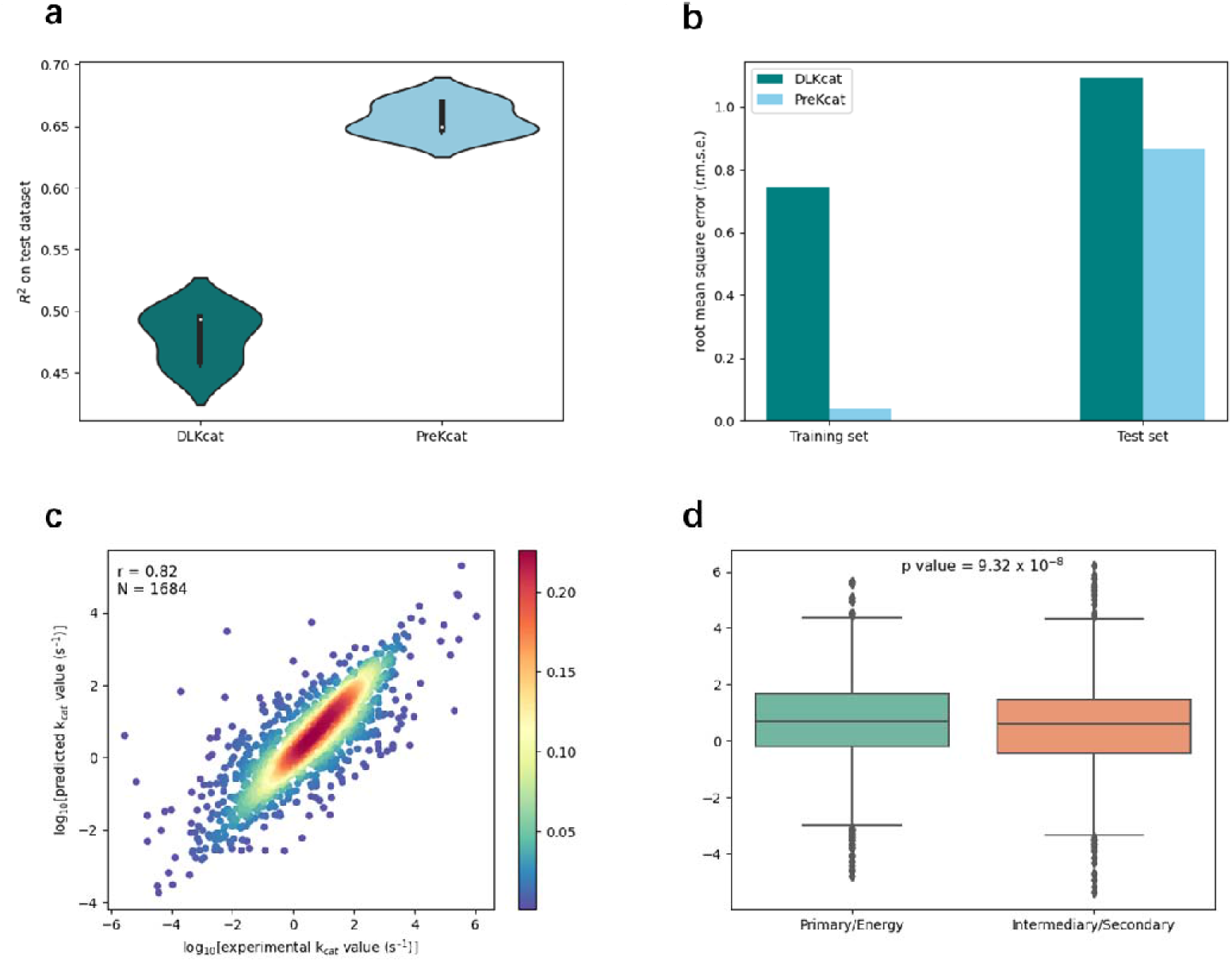
PreKcat prediction performance. a, Coefficients of determination of PreKcat and DLKcat on the test dataset by running the model five times. b, Root mean square error (r.m.s.e.) of PreKcat and DLKcat between predicted *k*_cat_ values and experimentally measured *k*_cat_ values on the training and test sets. c, Pearson correlation coefficient (p.c.c.) between the predicted *k*_cat_ values and the experimentally measured *k*_cat_ values on the test set. d, The *k*_cat_ value distribution of enzymes originating from primary central and energy metabolism and intermediary and secondary metabolism.

Furthermore, to prove that PreKcat could also discriminate the enzymes of different metabolic contexts, we divided all enzymes and their corresponding substrates into two categories according to DLKcat, which is primary central and energy metabolism, intermediary and secondary metabolism. Theoretically, the former should be higher than the latter. The results showed that the category of primary central and energy metabolism was significantly higher than the latter, which matches expectations (Fig. 2d; p = 9.32 x 10^-8^). In addition, PreKcat could also identify minor effects on enzyme variants with a few sites. The whole dataset was partitioned into wild-type and mutated enzymes. Great prediction performances were demonstrated on both wild-type enzymes (Fig. 3a for the test set; p.c.c. = 0.75; Supplementary Fig. 6 for the whole dataset; p.c.c. = 0.98) and mutant enzymes (Fig. 3b for the test set; p.c.c. = 0.89; Supplementary Fig. 7 for the whole dataset; p.c.c. = 0.99). The results were 10% and 11% higher than DLKcat on the test set for wild-type and mutant enzymes, respectively.

**Fig. 3.**
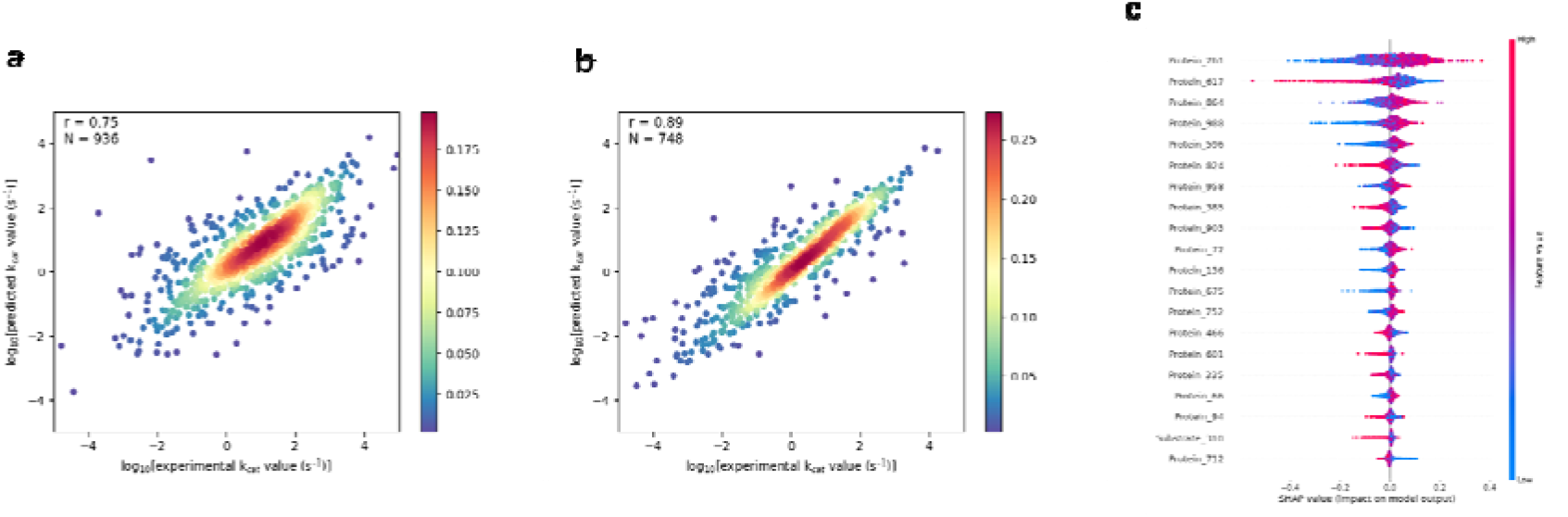
PreKcat prediction performance and feature importance analysis. a, Pearson correlation coefficient (p.c.c.) between predicted *k*_cat_ values and experimentally measured *k*_cat_ values of wild-type enzymes on the test set. b, Pearson correlation coefficient (p.c.c.) between predicted *k*_cat_ values and experimentally measured *k*_cat_ values of mutants on the test set. c, Distribution of SHAP values of every feature on the test set based on the trained PreKcat model.

To better understand feature preference during the model learning process, we employed SHapley Additive exPlanations (SHAP) to evaluate feature importance [22]. The higher the value is, the more important the feature. We calculated the SHAP value of every enzyme or substrate embedded feature on the test set based on the trained PreKcat model. The results showed that 19 of the top 20 features belonged to the enzyme category, while the rest belonged to the substrate (Fig. 3c). This confirmed that the embedded features of the enzyme had a more crucial effect than those of the substrate. In addition, for all of these top features, 10 of the top 20 features were positively correlated with the predicted *k*_cat_ values, while another 10 features were negatively correlated. The results showed that PreKcat has an obvious enzyme feature preference, meaning that enzyme information has a decisive effect.

### Weight redistribution improves *k*_cat_ prediction

To further solve real-world *k*_cat_ prediction problems effectively, i.e., those upon which biologists mainly and truly focus, such as high *k*_cat_ prediction and *k*_cat_ predictions of enzyme variants for specific substrates, we proposed a weight redistribution strategy to improve the results. This strategy relied on the adjustment of sample or feature weight to enhance the prediction ability on a specific issue (Fig. 1d).

Specifically, regarding the issue of high *k*_cat_ value prediction, a highly imbalanced dataset, with a few samples at both ends and most samples in the middle, which is close to a normal distribution (Fig. 4a), is difficult for a machine learning model to learn, resulting in a high error on high *k*_cat_ value prediction (Fig. 4b). To avoid this, we compared two representative sample weight optimization methods: directly modified sample weight (DM) and label distribution smoothing (LDS) [17]. We set experimentally measured *k*_cat_ values higher than 5 (logarithm value) as high values. This showed that the DM method is more effective (Supplementary Fig. 8). The weight of samples with high values was set to ten times that of other samples. This approach could effectively reduce the error in the high-value prediction, which demonstrated the superiority of the optimized model (Fig. 4c). Surprisingly, the overall prediction performance of PreKcat was basically unchanged (Fig. 4d). Altogether, by creating a sample weight redistribution strategy, we improved the prediction of high *k*_cat_ values.

**Fig. 4.**
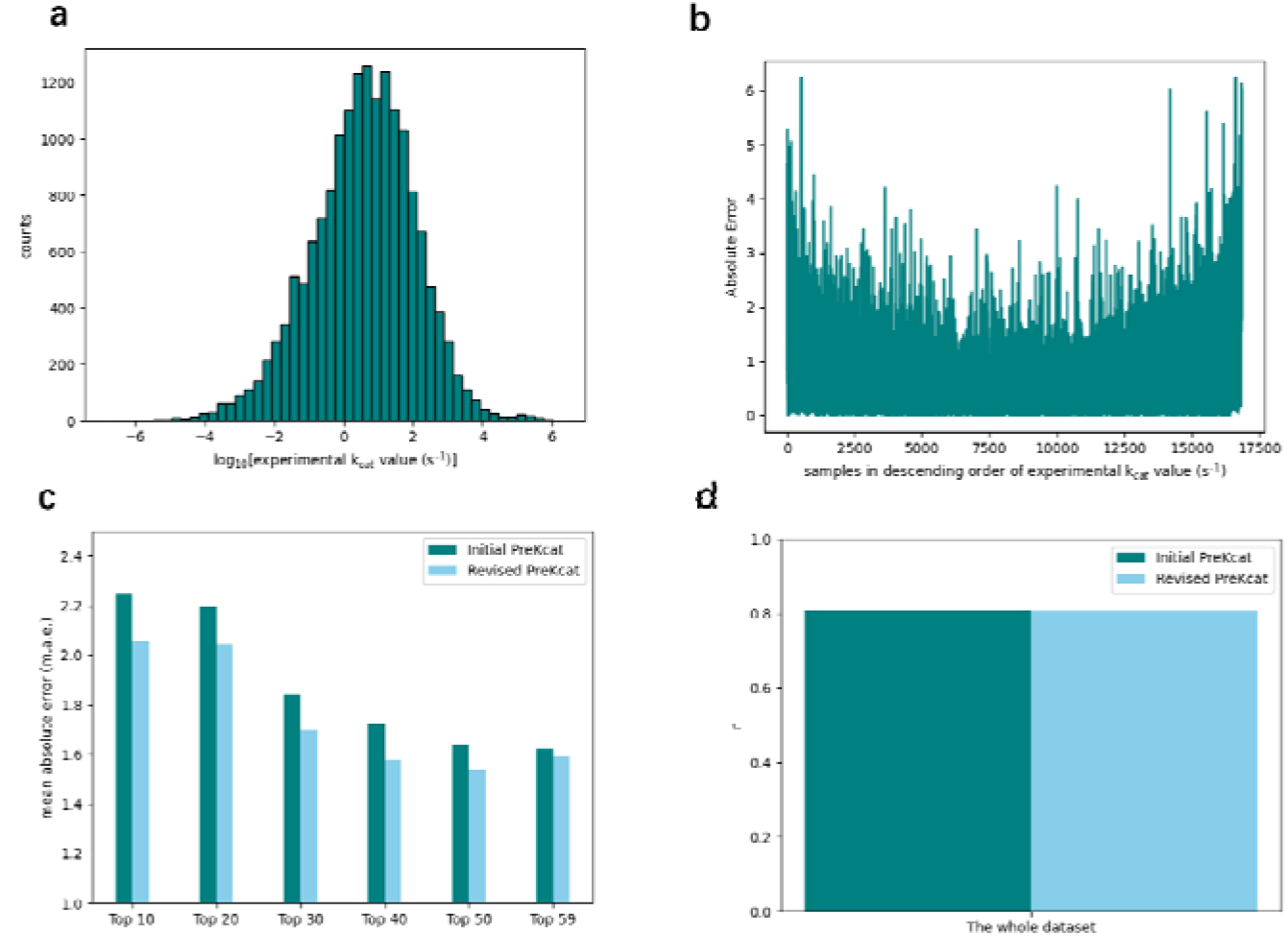
Sample weight redistribution for high *k*_cat_ prediction. a, Distribution of experimentally measured *k*_cat_ values. b, Predicted error distribution in descending order of experimentally measured *k*_cat_ values c, Mean absolute error (m.a.e.) of high *k*_cat_ value prediction. d, Pearson correlation coefficient (p.c.c.) of the initial and revised PreKcat on the whole dataset.

Weight redistribution also provided a feasible solution to address enzyme variant prediction problems. Due to the high importance of enzyme variant prediction, it has become a booming field. However, this issue related to mutants with a few mutated sites is usually not easy to predict, as is *k*_cat_. A previous prediction model made it difficult to differentiate the *k*_cat_ values of the wild-type enzyme and its mutants for a specific substrate if there were relatively few mutated sites or even just one (Supplementary Fig. 9). In addition, the initial PreKcat could not capture these minor variations because the embedded vectors only used comprehensive information (Supplementary Fig. 10). Therefore, we selected two crucial enzymes and their mutants in the naringenin synthetic pathway, tyrosine ammonia lyase (TAL) and chalcone synthase (CHS) (Fig. 5a). By setting a weighted pool strategy, embedded enzyme features could be used to pay more attention to mutated sites, leading to a significant distinction (Fig. 5b-d). As mentioned above, weight redistribution could provide an effective solution to identify minor amino acid substitutions.

**Fig. 5.**
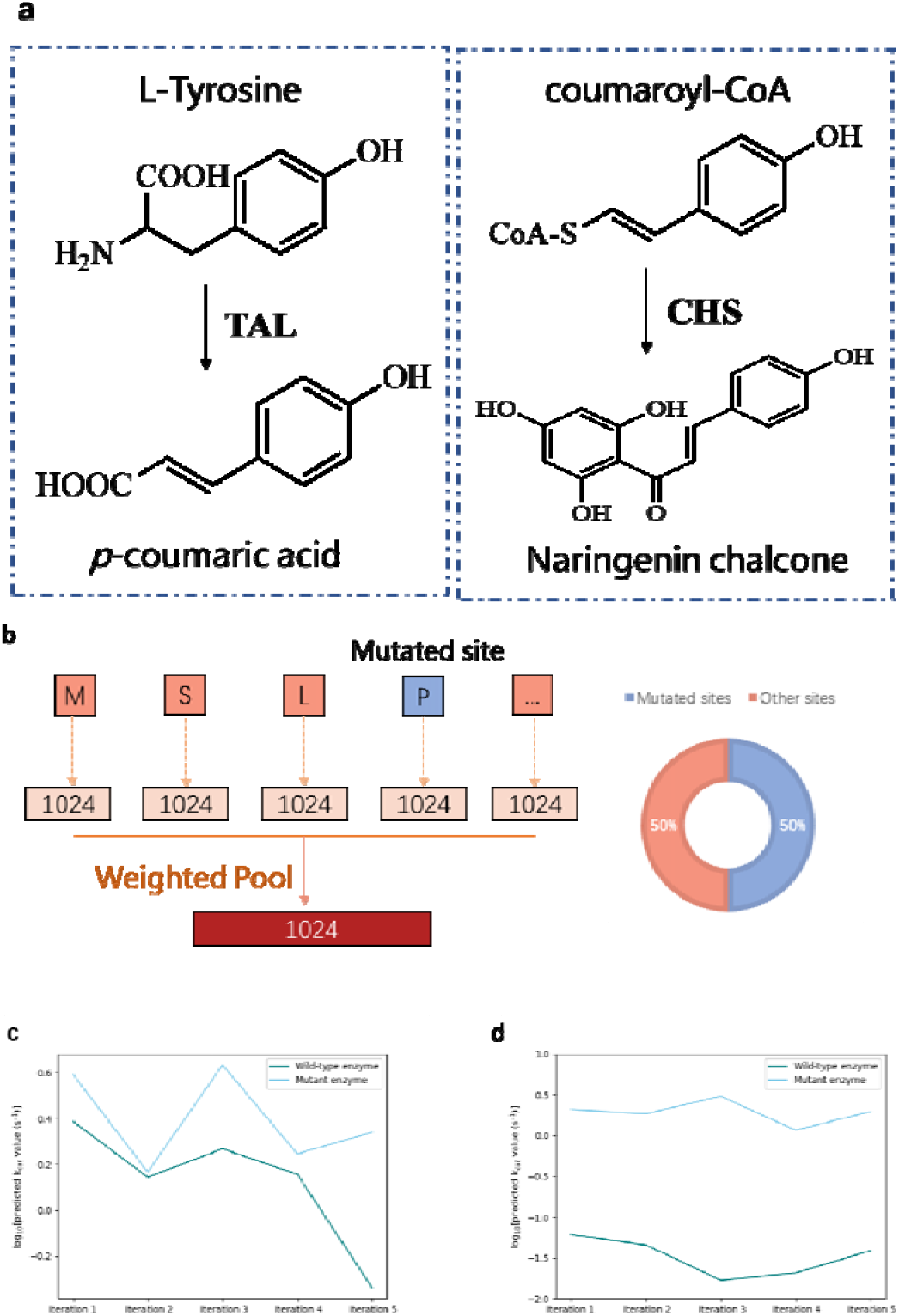
Feature weight redistribution for *k*_cat_ prediction. a, Reaction of two crucial enzymes on the naringenin synthetic pathway b, Feature weight redistribution strategy. c, Predicted *k*_cat_ values of wild-type TAL and its mutant by running the model five times. d, Predicted *k*_cat_ values of wild-type CHS and its mutant by running the model five times.

## Discussion

In this work, we proposed a pretrained language model-based Kcat prediction method (PreKcat), which achieved precise *k*_cat_ prediction solely from the enzyme sequence and substrate structure. We first proposed a framework based on representation and machine learning modules where the representation module relied on a pretrained protT5-XL and the SMILES Transformer to generate an effective representation, and the latter utilized a machine learning model to effectively capture the relationship between representation and *k*_cat_. Upon constructing this framework, we compared our prediction results with a state-of-the-art model, DLKcat. The average value of the coefficient of determination on the test dataset of five times of random splitting was 0.66, and the root mean square error, mean absolute error, and Pearson coefficient correlation all presented a remarkable advantage. We also verified that the performance on the test dataset where either the enzyme or substrate was not in the training set and on different metabolic contexts of the enzymes was advantageous. We demonstrated that PreKcat could predict different enzyme types, including wild-type and mutated enzymes. Furthermore, we sought to understand the feature preference during the model learning process, which concluded that protein information has a dominant effect.

Furthermore, we employed a weight redistribution strategy to adjust the sample weight for better prediction of high *k*_cat_ values. We compared different sample weight adjustment strategies and found that the DM strategy was most effective. Meanwhile, the revised model performs almost well with the initial PreKcat. In addition, we also utilized a weight redistribution strategy to adjust feature weights for tackling enzyme variant prediction, which realized a significant distinction for two crucial enzymes and their variants on the naringenin synthetic pathway. These results demonstrated that weight redistribution could help biologists make strong high *k*_cat_ value and enzyme variant predictions.

It is believed that PreKcat would provide a more powerful and useful tool for *k*_cat_ prediction, which would shed light on the development of enzyme kinetic parameters and downstream related tasks, such as enzyme evolution and drug design. Meanwhile, it is worth noting that this unified framework would also provide the possibility to precisely predict an enzyme’s Michaelis constant and protein-peptide interactions solely from protein sequences and molecular structures. Nevertheless, there are still some limitations to the current PreKcat. For instance, although PreKcat could differentiate some cases of experimentally measured enzymes and their variants, the predicted *k*_cat_ values were not sufficiently accurate. This may be caused by an insufficient dataset compared with known protein sequences and substrate structures. Additionally, the weight redistribution strategies could relieve the prediction bias caused by imbalanced datasets to some extent, but this has room to be further improved. An increasing number of experimentally measured *k*_cat_ values would also facilitate model performance. In the future, we plan to incorporate other algorithms that can effectively process imbalanced datasets, such as the synthetic minority oversampling technique, to further improve *k*_cat_ prediction.

## Methods

### Dataset source and preprocessing

We used the dataset provided by DLKcat, which was the most representative and largest dataset based on the enzyme sequences and substrate structures. The initially downloaded dataset comprised 17010 unique samples, and we excluded those samples whose substrate SMILES contained or *k*_cat_ values smaller than or equal to 0 referring to the DLKcat, resulting in 16838 samples. These samples contained 7822 unique protein sequences from 851 organisms and 2672 unique substrates. All *k*_cat_ values were converted into a logarithmic scale. Furthermore, to evaluate PreKcat, we randomly divided the whole dataset into training and test sets by 90% and 10% and repeated the process five times to evaluate the robustness of PreKcat.

### Construction of PreKcat

We implemented the PreKcat framework using torch v. 1.10.1+cu113 and sklearn v. 0.24.2. PreKcat consists of a representation and machine learning module. The representation module is responsible for generating an effective representation of the enzyme sequences and substrate structures. For the enzyme sequence, we used the representative pretrained protein language model protT5-XL, which has been verified to be effective on many peptide and protein function prediction tasks, to generate an embedded vector. Because previous studies have confirmed that mean pooling was most effective, every amino acid was converted into a 1024-dimensional vector on the last hidden layer, which was then summed and averaged. The final enzyme representation was an embedded 1024-dimensional vector. For the substrate structure, the simplified molecular-input line-entry system (SMILES) was generated and imputed into the pretrained SMILES transformer, which has been verified to have great performance on various molecular property prediction tasks. It would also generate a 1024-dimensional vector for the substrate, which contains the mean and max pools of the penultimate and last hidden layers. By an unsupervised learning process, the enzyme sequence or substrate structure would be converted into a numerical representation rather than a series of chars initially, which is hard for machine learning models to learn. The second module was the Extra Tree model, a representative machine learning method that can effectively capture the relationship between concatenated representation vectors of enzyme sequence and substrate structure and the *k*_cat_ value. All experiments were tested in a Linux environment running Ubuntu 20.04.5 on a server with 64 cores and 4 NVIDIA GeForce RTX 3080 GPUs. Only a single core and a GPU were used for training.

### Evaluation metrics

To evaluate model performance, we considered various evaluation metrics for predicting the *k*_cat_ value and the experimentally measured *k*_cat_. Specifically, we selected the coefficient of determination (R^2^) in Equation 1, the Pearson correlation coefficient (p.c.c.) in Equation 2, the root mean square error (r.m.s.e.) in Equation 3, and the mean absolute error (m.a.e.) in Equation 4, where *y_ie_* denotes the experimentally measured *k*_cat_ value, *y_ip_* denotes the predicted *k*_cat_ value, 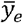 denotes the average of the experimentally measured *k*_cat_ values, 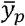 denotes the average of the predicted *k*_cat_ values, and n denotes the number of samples (depending on the size of the selected dataset).

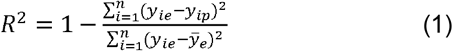

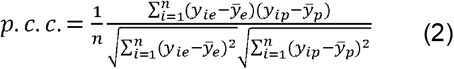

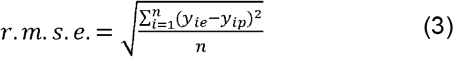

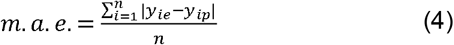

### Feature importance analysis by SHAP

We employed SHapley Additive exPlanations (SHAP), a unified framework for model interpretability analysis, which could calculate an importance value for each feature. The higher the assigned SHAP value is, the more important the feature. Additionally, it can reflect the positive or negative effects of features. It has been used to interpret the importance of various biological problems, such as Type IV Secreted Effectors prediction and anticancer peptide prediction. Therefore, we tested it on the *k*_cat_ test dataset for accelerating calculation time, which contains 1684 samples, based on the trained PreKcat model. The SHAP summary from TreeExplainer presented the magnitude, distribution, and direction of every feature effect. Each dot represents a sample of the dataset. The position along the x-axis denotes the SHAP value, and the change in colour denotes different feature values. The SHAP had a freely available Python package and was implemented here.

### t-distributed stochastic neighbour embedding (t-SNE) visualization

To understand the distribution of embedded enzyme or substrate representation and to explore the necessity of following machine learning, we utilized t-SNE to visualize the embedded enzyme and substrate vectors. This method has been widely used to analyse the feature distribution of biological tasks, such as antimicrobial peptide recognition and protein subcellular location. We selected all 16838 samples and calculated their embedded vectors. The concatenated vectors were then imputed into the t-SNE and converted into a 2-dimensional vector for each sample. The default parameters were set automatically. Then, for the convenience of display, we normalized the calculated numerical values of the 2-dimensional vector in Equation 5, where *y_it_* denotes the value of the i_th_ projected vector and *y_t_* denotes all values of the projected vector.

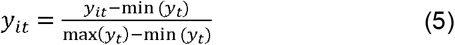

### Sample weight redistribution strategies

We used representative weight redistribution strategies to adjust the weight of the sample for precise high *k*_cat_ prediction. Specifically, for sample weight adjustment, we compared two representative sample weight optimization methods: directly modified sample weight (DM) and label distribution smoothing (LDS). DM means that the weight of samples with *k*_cat_ values higher than 5 (logarithm value) would be enhanced. We compared several parameters, including the weight multipliers (5, 10, 20, 50) and whether they were normalized. This resulted in eight optimized model combinations. These results were all provided (Supplementary Fig. 8). LDS is a simple, effective, and interpretable algorithm for tackling the problem of unbalanced datasets that exploits the similarity of the nearby label space. It had been verified to be very effective in sections where only a few samples exist, and the predicted error would be reduced dramatically. LDS convolves a symmetric kernel with the empirical density distribution to generate a kernel-smoothed effective density distribution in Equation 6, where *p*(*y*) denotes the number of appearances of label *y* in the training data, 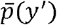 denotes the effective density of label *y*′, and *k*(*y,y′*) denotes the symmetric kernel. Similarly, we selected a Gaussian kernel and set various kernel sizes (3, 5, 7) and sigma values (1, 2). For the convenience of comparison, we employed 5-fold cross-validation on the whole dataset. The predicted *k*_cat_ values were divided into different intervals to calculate the error (top 10, 20, 30, 40, and 50).

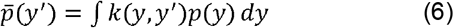

### Feature weight redistribution strategies

We also implemented feature weight redistribution strategies to optimize the results of enzyme variants with a few mutated sites. To capture the minor effects of mutated sites, we abandoned the mean-pooling strategy to generate a per-protein representation of the enzyme. Instead, we assigned one half of a whole weight pool to the mutated amino acid site, while the rest shared the remainder. The value was calculated using Equation 7, where *w_i_*, denotes the weight of the i_th_ mutated site and S denotes the number of mutated sites, as do other unmutated sites.

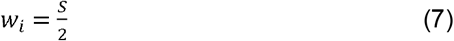

### Experimentally measured *k*_cat_ of enzyme variants

To validate the effectiveness of feature weight redistribution, we selected two unpublished crucial enzymes on the naringenin synthetic pathway and their mutants with only one site. The tyrosine ammonia lyase (TAL) and chalcone synthase (CHS) enzymes are crucial in the naringenin synthetic pathway (Fig. 5a). The original and mutated sequences are both provided in the Supplementary information. The mutated site of TAL was 174, at which histidine was converted into glutamine. The mutated site of CHS was 293, at which serine was converted into threonine. The real values of *k*_cat_ of the wild-type and mutant enzymes were experimentally measured.

## Data availability

All data analysed in this article are included in this manuscript and supplementary information.

## REFERENCES

1. Chen, Y. & Nielsen, J. Energy metabolism controls phenotypes by protein efficiency and allocation. Proc. Natl Acad. Sci. USA 116, 17592–17597 (2019).

2. Sánchez, B. J. et al. Improving the phenotype predictions of a yeast genomescale metabolic model by incorporating enzymatic constraints. Mol. Syst. Biol. 13, 935 (2017).

3. Adadi, R., Volkmer, B., Milo, R., Heinemann, M. & Shlomi, T. Prediction of microbial growth rate versus biomass yield by a metabolic network with kinetic parameters. PLoS Comput. Biol. 8, e1002575–e1002575 (2012).

4. Klumpp, S., Scott, M., Pedersen, S. & Hwa, T Molecular crowding limits translation and cell growth. Proc. Natl Acad. Sci. USA 110, 16754–16759 (2013).

5. Apweiler, Rolf, et al. “UniProt: the universal protein knowledgebase.” Nucleic acids research 32.suppl_1 (2004): D115–D119.

6. Schomburg, I. et al. The BRENDA enzyme information system–from a database to an expert system. J. Biotechnol. 261, 194–206 (2017).

7. Wittig, U., Rey, M., Weidemann, A., Kania, R. & Müller, W. SABIO-RK: an updated resource for manually curated biochemical reaction kinetics. Nucleic Acids Res. 46, D656–D660 (2018).

8. Borger, Simon, Wolfram Liebermeister, and Edda Klipp. “Prediction of enzyme kinetic parameters based on statistical learning.” Genome Informatics 17.1 (2006): 80–87.

9. Heckmann, David, et al. “Machine learning applied to enzyme turnover numbers reveals protein structural correlates and improves metabolic models.” Nature communications 9.1 (2018): 1–10.

10. Li, Feiran, et al. “Deep learning-based *k*_ca_t prediction enables improved enzyme-constrained model reconstruction.” Nature Catalysis 5.8 (2022): 662–672.

11. Vaswani, Ashish, et al. “Attention is all you need.” Advances in neural information processing systems 30 (2017).

12. Devlin, Jacob, et al. “Bert: Pre-training of deep bidirectional transformers for language understanding.” arXiv preprint arXiv:1810.04805 (2018).

13. Elnaggar, Ahmed, et al. “ProtTrans: towards cracking the language of Life’s code through self-supervised deep learning and high performance computing.” arXiv preprint arXiv:2007.06225 (2020).

14. Rives, Alexander, et al. “Biological structure and function emerge from scaling unsupervised learning to 250 million protein sequences.” Proceedings of the National Academy of Sciences 118.15 (2021): e2016239118.

15. Honda, Shion, Shoi Shi, and Hiroki R. Ueda. “Smiles transformer: Pre-trained molecular fingerprint for low data drug discovery.” arXiv preprint arXiv:1911.04738 (2019).

16. Chawla, Nitesh V. “Data mining for imbalanced datasets: An overview.” Data mining and knowledge discovery handbook (2009): 875–886.

17. Yang, Yuzhe, et al. “Delving into deep imbalanced regression.” International Conference on Machine Learning. PMLR, 2021.

18. Anand, Ashish, et al. “An approach for classification of highly imbalanced data using weighting and undersampling.” Amino acids 39.5 (2010): 138561391.

19. Agrawal, Piyush, et al. “AntiCP 2.0: an updated model for predicting anticancer peptides.” Briefings in bioinformatics 22.3 (2021): bbaa153.

20. Agrawal, Saurabh, Dilip Singh Sisodia, and Naresh Kumar Nagwani. “Multi-function prediction of unknown protein sequences using multilabel classifiers and augmented sequence features.” Iranian Journal of Science and Technology, Transactions A: Science 45.4 (2021): 1177–1189.

21. Van der Maaten, Laurens, and Geoffrey Hinton. “Visualizing data using t-SNE.” Journal of machine learning research 9.11 (2008).

22. Lundberg, Scott M., and Su-In Lee. “A unified approach to interpreting model predictions.” Advances in neural information processing systems 30 (2017).

